# Histone deacetylase inhibitor induces acetyl-CoA depletion leading to lethal metabolic stress in RAS-pathway activated cells

**DOI:** 10.1101/2022.01.31.478570

**Authors:** Agnes Basseville, Pierre-Christian Violet, Maryam Safari, Carole Sourbier, W. Marston Linehan, Robert W. Robey, Mark Levine, Dan L. Sackett, Susan E. Bates

## Abstract

RAS-mutant cancers are among the most refractory to treatment. Apart from new G12C genotype targeted therapies, strategies to kill RAS-mutant cells by directly targeting RAS or its downstream effectors have been mostly unsuccessful, mainly due to pathway redundancy and heterogeneities in RAS-induced phenotypes. Here we identified a RAS-phenotype that can be targeted by the histone deacetylase inhibitor (HDACi) romidepsin. We showed that the hyperacetylation induced by romidepsin depleted acetyl-CoA, the cell donor substrate for acetylation, and led to metabolic stress and death in KRAS-activated cells. Elastic net analysis on transcriptomics from a 608-cell panel confirmed that HDACi sensitivity was linked to a difference in profiles in two pathways involved in acetyl-CoA metabolism. The analysis of a clinical dataset confirmed that perturbation of the two acetyl-CoA pathways were correlated with HDACi sensitivity in patients treated with belinostat. Our analysis suggests the potential utility of a RAS-associated acetyl-CoA phenotype to sharpen treatment choices for RAS-activated tumors.

## INTRODUCTION

RAS-mutated tumors are the most common and the deadliest of all human cancers, and remain among the most refractory to available treatments (1). Accordingly, the highest prevalence of RAS mutation is found in cancers with the lowest 5-year survival rates: pancreatic cancer (9% survival), non-small cell lung cancer (23%) and colorectal cancer (64%) (2). In the wild-type (WT) state, RAS protein activation/deactivation controls central cellular processes such as growth, migration or survival, as well as interactions with cells from the extracellular environment like immune cells. When mutated, RAS remains in its active state and continuously activates its downstream effectors (RAF, PI3K, RALGDS, TIAM1) leading to aberrant signaling in these pathways (1).

Multiple therapeutic strategies have been designed to shut down RAS pathways by targeting the more common form of RAS (KRAS) directly, or its most common downstream effectors (for instance RAF/MEK or PI3K/Akt). Several inhibitors have reached the clinic, but challenges remain to reach the breadth of RAS mutations, and to deepen treatment responses (1,3,4). The main difficulties encountered in these efforts are the compensation by redundant paralogs, the redundancies in RAS-activated pathways and the existing feedback loops (5,6). Also, some tumors are described as RAS-addicted, but others maintain viability despite RAS inhibition, and this RAS dependency has been shown to vary widely even within the same cancer type (7,8).

One lead that has emerged involves the exploitation of tumor metabolic vulnerabilities induced by RAS activation (9). In fact, cancer cells rewire their metabolic pathways to meet their needs in energy and biosynthetic precursors, both essential for unrestrained proliferation. For example, in pancreatic cancer, KRAS promotes a shift in the glutamine pathway to support redox maintenance, as well as an increase in glycolysis to divert nutrients into the glycosylation and pentose phosphate pathways. KRAS-driven cancers also recycle metabolites through increased autophagy or macropinocytosis to meet their increased demand for metabolites. Identification of these KRAS-tumor-specific and potentially targetable metabolic pathways led to new clinical trials involving pyruvate dehydrogenase inhibition in patients with KRAS-mutated pancreatic cancers, and glutaminase inhibition in lung cancer (NCT03699319, NCT02071862). Importantly, these KRAS-linked metabolic reprograming features differ according to KRAS copy number, tumor types, and genetic context (10,11). In accordance, scientists pinpointed the development of biomarkers as critical to allow selection among patients bearing oncogene-driven tumors of those more likely to benefit from metabolic-targeting drugs (12).

Interestingly, our group and others (13–15) have shown that RAS-transformed cells display enhanced sensitivity to the histone deacetylase (HDAC) class I inhibitor romidepsin, although the mechanism of sensitivity has not yet been elucidated. HDAC inhibitors (HDACis) are epigenetic modulators initially known to induce hyperacetylation of histone proteins leading to transcriptional deregulation. Other biological consequences of HDAC inhibitors have also been demonstrated, including accumulation of RNA-DNA hybrids known as R-loops, DNA damage, oxidative stress, cell cycle arrest and apoptosis (16,17). It was approved by the U.S. Food and Drug Administration in 2009 to treat T-cell lymphoma but has limited effect in monotherapy for solid tumors. Its pleiotropic mechanism of action remains of high interest in the scientific community in finding appropriate drug combinations for solid tumors in clinic (16).

In this study, we show that romidepsin-induced hyperacetylation caused depletion of stores of acetyl-CoA, the cell donor substrate for acetylation, and leads to lethal metabolic stress in KRAS-activated cells but not in KRAS-WT. We also confirmed that two acetyl-CoA metabolism pathways are hallmark of sensitivity in a cell panel and in patients treated with an HDACi, validating not only these drugs as metabolism-targeting drugs, but also these pathways as a potential tool to identify a RAS-linked metabolic phenotype in a personalized medicine strategy.

## MATERIALS and METHODS

### Cell lines, treatment and statistical tests

The complete NCI-60 panel was obtained from National Cancer Institute (NCI) Anticancer Drug Screen (Bethesda, MD). Pancreatic cell lines were obtained from ATCC (Manassas, VA). All cells were cultivated in RPMI including 2mM glutamine and 11.11mM glucose supplemented by 10% FBS and penicillin/streptavidin (GIBCO, Grand Island, NY).

Romidepsin (depsipeptide, NSC 630176) was provided by the Cancer Therapy Evaluation Program, NCI. Tariquidar was provided by the NCI Anticancer Drug Screen. PD-0325901 (MEK inhibitor), MK-2206 (AKT inhibitor) were purchased from ChemieTek (Indianapolis, IN). Glutaminase inhibitor 968 compound was obtained from Millipore (Burlington, MA); oligomycinA, NAC, glutathione, dimethyl-α-ketoglutarate, dimethyl-glutamate, sodium citrate and sodium acetate from Sigma-Aldrich (St. Louis, MO); DPI, ML171 and etomoxir from Selleckchem (Houston, TX). MitoQ was provided by Dan L. Sackett (NIH, NICHHD).

Unless otherwise specified, t-tests were performed to evaluate significance of results, and asterisks were added to graphs when p-value was lower than 0.05.

### Transcriptomic analysis with R

Gene expression basal level for NCI-60 and CCLE panel were downloaded via the cBio Cancer Genomics Portal (18). Gene expression for acute myeloid leukemia samples was downloaded from Gene Expression Omnibus (dataset GSE39363). Response to eight HDACi for CCLE panel were obtained from The Genomics of Drug Sensitivity in Cancer Project public database (19). Gene expression for romidepsin-treated cells was obtained as follows. Cells were harvested after treatment and RNA extraction was performed using RNeasy Mini Kit (QIAGEN, Hilden, Germany), following manufacturer’s instruction. Samples in triplicate were sent to the Frederick Cancer Research Core (NIH, NCI), where mRNA levels were determined by using Affymetrix GeneChip® Human Genome U133 Plus 2.0 Array. Microarray data have been deposited in NCBI Gene Expression Omnibus (accession number GSE133120). Detailed methods are provided in related figure legends.

### Mouse study

Animal procedures were approved by the Animal Care and Use Committees at NIH/NIDDK. Athymic female mice (Ncr-nu/nu, 5-6 weeks old, ∼20g) were purchased from The Jackson Laboratory (JAX, Maine, USA). Tumors were generated by injecting 10 million cells subcutaneously into left flank. Treatment started once tumor size reached an average of 100mm^3^. Mice received 3mg/kg romidepsin by intraperitoneal injection every 4 days and/or 5mg/kg PD-0325901 by oral gavage every 4 days. Procedures are detailed in Supplementary Figure 1.

### Flow cytometry

Cell death was measured by flow cytometer after annexin V-FITC/ propidium iodide staining, according to manufacturer’s instruction (eBioscience, Inc., San Diego, CA). Cell lines with high expression of the multidrug efflux transporter P-glycoprotein (P-gp) according to cellMiner database (20) were co-treated with a Pgp inhibitor to avoid the bias of drug efflux during the assessment of toxicity. Reactive Oxygen Species (ROS) quantification was performed with 10µM H2DCF-DA (Abcam, Cambridge, UK) or 1µM MitoSOX™ Red (Invitrogen, Carlsbad,CA). 1µM tariquidar and 50µM MK571 (Selleckchem) were added to medium to inhibit fluorescent compound efflux by ABC transporters. Mitochondrial membrane potential was measured after incubation of cells in phenol red-free medium with 5µM JC1 (Invitrogen) and 1µM tariquidar. Cytochrome c release was measured by flow cytometry using the Cytochrome c Release Apoptosis Assay Kit (Millipore).

### Western blot

Western blot was performed as previously described (13). Histone H3 antibodies were purchased from Cell Signaling Technology (Danvers, MA), acetyl-histone H3 from Upstate Biotechnology (Lake Placid, NY), γH2AX from Millipore, and GAPDH from American Research Products (Waltham, MA).

### Seahorse experiment

Oxygen consumption rates (OCR) and extracellular acidification rates (ECAR) were measured using the XF96 or XFp Extracellular Flux analyzer (Seahorse Bioscience, North Billerica, MA). Protein amount per well was quantified using BCA assay.

### Metabolomics analysis

Metabolic profiling of A549 and ACHN cell lines was performed by Metabolon Inc. (Durham, NC). After treatment, cells were pelleted, frozen and sent for analysis (quadruplicates). All samples were randomly distributed across a single day platform run on GC/MS and LC/MS/MS instrumentation, and they were analyzed using Welch’s two-sample T-test.

### Acetyl-CoA quantification analysis

Cells were harvested and sent frozen to the Metabolomics Core at the Translational Research Institute for Metabolism and Diabetes at Florida Hospital for estimation of acetyl-CoA and malonyl-CoA by LC-MS/MS.

## RESULTS

### Activated RAS pathway is linked to romidepsin sensitivity

To identify the determinants of sensitivity to romidepsin alone or in combination with RAS downstream inhibitors (MEK and AKT inhibitors), we screened the NCI-60 panel (58 human cell lines from seven cancer tissues), adding seven pancreatic cancer cell lines because they have a high prevalence of RAS mutation (97%) (1). For this and all the experiments in this article, romidepsin was used with a dosing mimicking its concentration measured in the clinic, referred as “clinical dosing” (46nM romidepsin, with drug removal at 6h when treatment time was over 6h (21)). All the other treatments and co-treatments were added continuously for the indicated concentrations. The 65 cells were hierarchically clustered according to their response to treatment (Figure 1a), with four major clusters formed, mainly based on response to romidepsin alone. The addition of RAS downstream inhibitors increased romidepsin efficacy proportionally (Figure 1b), with a 0.87 correlation in the 65 cell lines (p-value>0.0001) between romidepsin alone and romidepsin in combination with MEK inhibitor (MEKi, PD-0325901), and a 0.92 correlation (p-value>0.0001) for the same analysis with AKT inhibitor (AKTi, MK-2206). All 65 lines but two were insensitive to AKTi or MEKi alone. These results indicate that RAS downstream inhibitors did not trigger cell death, but amplified romidepsin-induced cell death. We queried mutations in well-known key cancer genes carried by these cell lines from the Cosmic Cancer Database. Eleven relevant genes were grouped in four categories: TKR/RAS/BRAF pathway (EGFR, Erb2, KRAS, NRAS, HRAS and BRAF), PI3K pathway (PTEN, PI3CA and PI3R1), CDKN2A, and p53 (Figure 1a). Noticeably, the cluster of cells that did not respond to treatment correlated with absence of mutation in TKR/RAS/BRAF pathway, with a 0.47 correlation (p-value=0.002). Cell doubling time, mutations in the PI3K pathway, CDKN2A or p53 did not correlate with treatment efficacy (Supplementary Figure 1a-b). Basal expression levels for the 65-cell panel (18) were analyzed by a Significance Analysis of Microarrays (SAM) to identify significant genes in romidepsin non-sensitive and sensitive clusters. The significant genes were analyzed by Gene Set Enrichment Analysis (GSEA) software using the hallmark gene sets. Confirming the mutation analysis, “KRAS signaling up” gene signature was significantly enriched (nominal p-value<0.01 and FDR q-value<0.05) in the two romidepsin-sensitive cell clusters (Figure 1c). Gene expression value of the leading-edge genes from the signature in cluster 2 and 4 were plotted on a heatmap (Figure 1d). Again, cell lines resistant to romidepsin clustered together based on decreased “KRAS signaling up” signature expression. The heatmap also illustrated the variability in KRAS signature-related genes according to organ site. Together, these results showed an association between RTK/RAS/BRAF pathway activation and romidepsin sensitivity.

**Figure 1:**
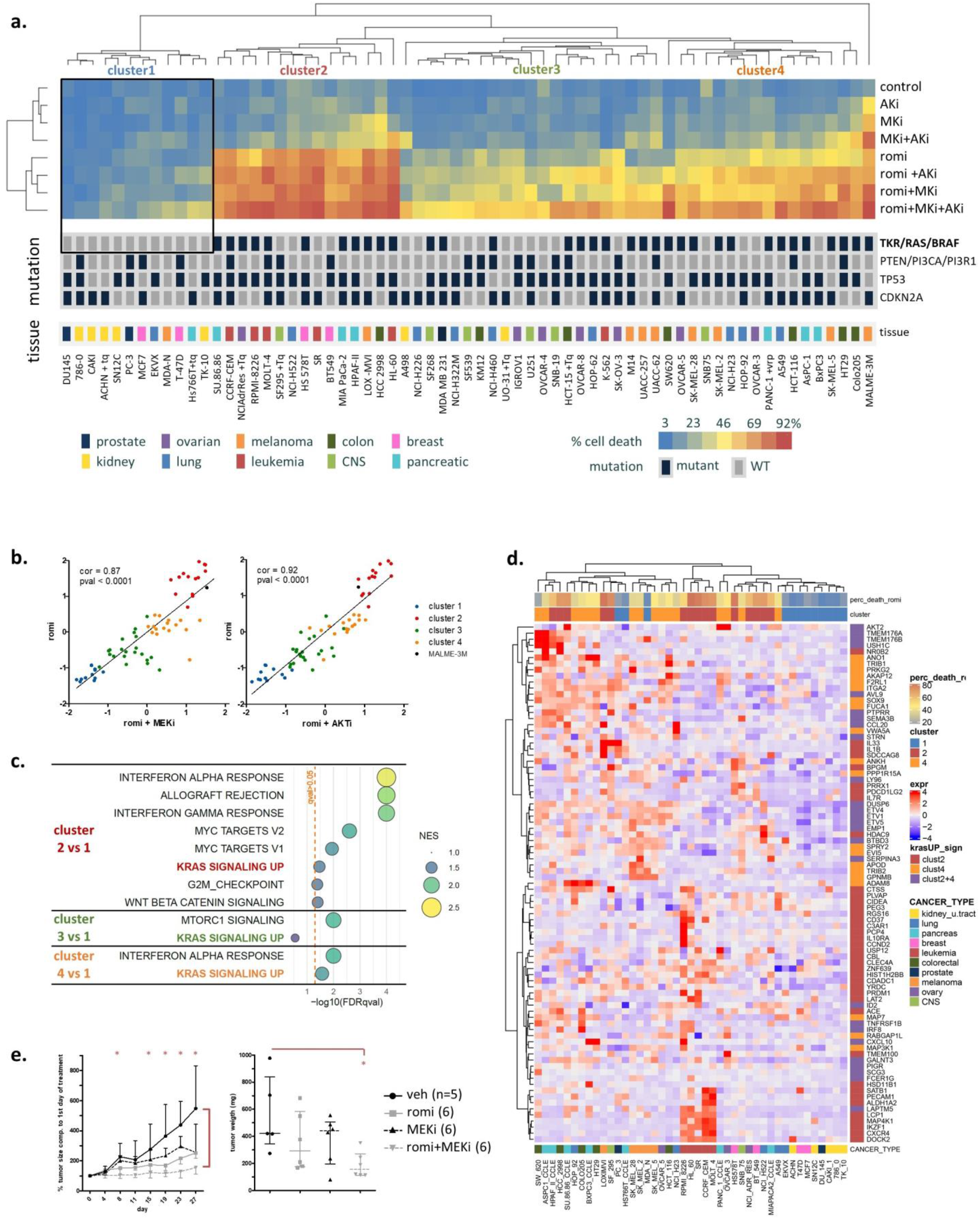
Activated RAS pathway is linked to romidepsin sensitivity. **a**. Cells were treated with romidepsin (clinical dosing) and/or MEK inhibitor (MEKi, 250nM) and/or AKT inhibitor (AKTi, 1µM), and cell death was assessed at 48h by annexin V-PI assay. Cell lines with high expression of the multidrug efflux transporter P-glycoprotein were cotreated with P-gp inhibitor Tariquidar (Tq) to avoid the bias of drug efflux during the assessment of toxicity. Cells were hierarchically clustered according to percentage cell death, and a heatmap was drawn accordingly. Key DNA mutations and tissue origin of cancer are indicated below the heatmap. **b**. The response to romidepsin in combination with either MEKi (left) or AKTi (right) for the 65 cell lines were plotted according to response to romidepsin alone (z-score). Cells were colored according to the four clusters obtained in Figure 1a. **c**. SAM analysis was performed on NCI-60 panel gene expression between low responsive cluster (cluster 1) and each of the three others. SAM-ranked genes were analyzed with GSEA, and the significant enriched gene sets (FDR q-value <0.05) obtained for each cluster was plotted according to normalized enrichment score (NES) and false discovery rate (FDR). **d**. Leading-edge genes for “KRAS up signaling” signature obtained by GSEA analysis in cluster 2 and 4 were hierarchically clustered according to their basal expression in the 65 cell lines. **e**. Athymic mice were injected with KRAS mutation bearing A549 cells forming tumor, then treated with romidepsin and/or MEKi for 28 days. Tumor size (median+/-IQR) and mice weight (Supplementary Figure 2c) were measured over time, and tumor weight (median+/-IQR) was measured after mice sacrifice and tumor dissection. Statistical difference between control and treated groups was determined using Welch’s t-test (unequal variance t-test).

To assess the efficacy of the combination drug therapy in a preclinical model, we injected A549 cells (KRAS-mutant, cluster 4) subcutaneously in athymic mice and treated them with romidepsin and/or MEKi (Figures 1e and Supplementary 1c). From 15 days of treatment, the tumors treated with romidepsin/MEKi were significantly smaller than the untreated ones. Tumor weight at sacrifice was significantly lower in mice treated with the combination compared to untreated mice whereas no significant difference was noticed after single agent treatment, confirming the efficacy of the combination *in vivo*.

### Early cytoplasmic ROS release differentiates KRAS sensitivity to romidepsin, while acetylation and DNA damage do not

To understand why RTK/RAS/BRAF mutated cells were sensitive to romidepsin, we focused on the KRAS-mutant cells as a model. Progression in signaling events during romidepsin treatment using clinical dosing was evaluated in at least three KRAS-mutant and three KRAS-WT cell lines.

Histone acetylation, the main target of romidepsin, and DNA damage (using the commonly accepted marker γH2AX) were measured by western blot after 6h combination treatment in six cell lines (Figure 2a). No differences were observed between KRAS-mutant and KRAS-WT cells.

**Figure 2:**
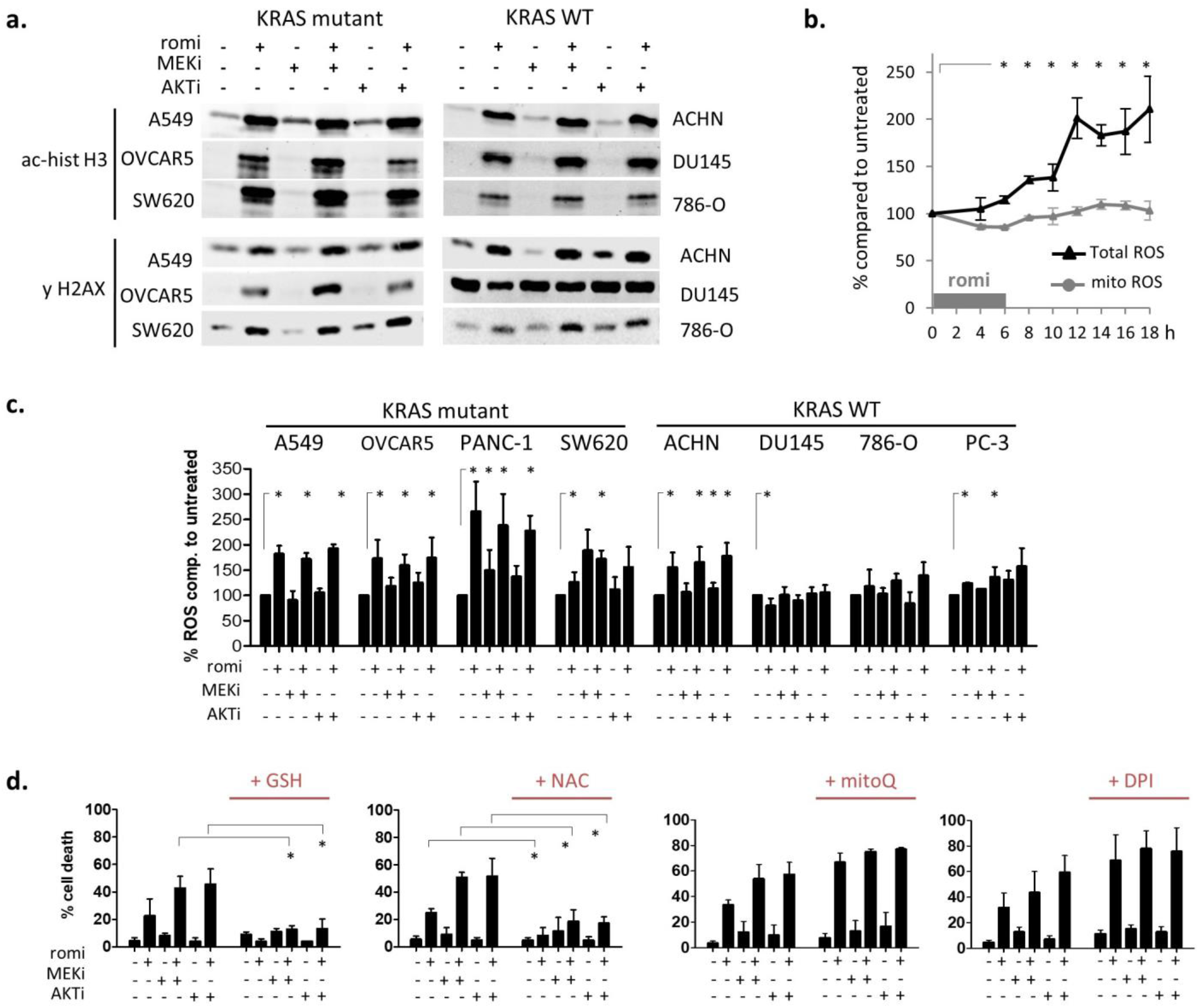
Early cytoplasmic ROS release differentiates KRAS sensitivity to romidepsin, while acetylation and DNA damage do not. **a**. Histone acetylation and H2AX phosphorylation (DNA damage marker) were measured by WB in cells treated for 6h with romidepsin +/- MEKi or AKTi. **b**. Total and mitochondrial ROS production were measured over time by flow cytometry for 18h during treatment with romidepsin (clinical dosing) in A549 cell line. **c**. Total ROS production was measured at 24h in cell lines after romidepsin (clinical dosing) +/- MEKi +/- AKTi. **d**. Percentage cell death was measured at 48h in A549 cell line after treatment with romidepsin +/- MEKi +/- AKTi, in presence or not of cytoplasmic ROS scavengers (5mM GSH or NAC), mitochondrial ROS scavenger (1µM MitoQ), or pan-NADPH oxidase inhibitor (5µM DPI). In b-d, data are plotted as mean of three experiments +/-SD, and statistical significance between groups was assessed using unpaired two-tailed Student’s t-test.

Both sensitive and resistant cells undergo increased histone acetylation when treated with romidepsin, and five out of six cell lines had increased DNA damage. MEKi and AKTi did not increase these signals.

Next, total and mitochondrial reactive oxygen species (ROS) production were measured over time in A549 cell line treated with romidepsin by flow cytometry. Total ROS increased around 10-12h, whereas no increase was detected in mitochondrial ROS over 18h (Figure 2b). Total ROS production was then evaluated in eight cell lines after 24h combination treatment (Figure 2c). ROS production was mainly induced in romidepsin-sensitive KRAS-mutant cells: only one out of four KRAS-WT cells had a significant romidepsin-induced ROS increase, whereas all four KRAS-mutant cells showed an increase. MEKi and AKTi did not increase ROS signaling.

ROS involvement in cell death at 48h was evaluated after the 6h romidepsin treatment. Scavenging of total ROS by glutathione (GSH) or N-acetyl-cysteine (NAC) significantly rescued all KRAS-mutant cells from death, whereas neither mitochondrial ROS scavenger (mitoQ) nor NADPH oxidase (DPI) inhibitor did (Figures 2d and Supplementary 2a). We also assessed mitochondrial ROS production, mitochondrial membrane depolarization and the apoptotic marker cytochrome c release at 24h and 48h (Supplementary Figure 2b). These signals - indicating onset of cell death - were low at 24h but high at 48h in romidepsin-treated KRAS-mutant cells. Combination with MEKi or AKTi amplified most of these late death signals at 48h.

Taken together, these results indicated that romidepsin induces early cytoplasmic ROS production in KRAS-mutant cells. Mitochondrial ROS production and mitochondrial membrane depolarization appeared late, suggesting that they were apoptotic by-products rather than drivers of apoptosis. Having eliminated the main known causes of ROS production (NADPH oxidase and mitochondria), we therefore explored other possible sources of cytoplasmic ROS imbalance after romidepsin treatment.

### Romidepsin affects two TCA-fueling pathways: fatty acid beta-oxidation and glutaminolysis

Four enzymes in metabolic pathways produce cytoplasmic NADPH, an intrinsic ROS scavenger that maintains cytoplasmic redox balance (Figure 3a). We hypothesized that romidepsin might affect metabolic pathways that regulate NADPH production and therefore ROS sensitivity. To investigate, we explored the metabolic effect of romidepsin by measuring in real time cell oxygen consumption rates (OCR), a surrogate of oxidative phosphorylation (OXPHOS), and cell extracellular acidification rates (ECAR), a surrogate of glycolysis. We first assessed basal OCR and ECAR in nine cell lines treated for 18h with romidepsin using clinical dosing (Figure 3b). Romidepsin treatment decreased OXPHOS in the four KRAS-mutant cells, and in two out of five KRAS-WT cells. Next, by varying fatty acid, glucose, or glutamine supply, we could evaluate the three main tricarboxylic acid (TCA) cycle fueling pathways: fatty acid beta-oxidation, glycolysis, and glutaminolysis. We evaluated fatty acid β-oxidation-supported OXPHOS by adding etomoxir, an inhibitor of carnitine palmitoyltransferase-1, rate-limiting-enzyme in β-oxidation. Etomoxir decreased OCR for all untreated cell lines, indicating that all cell lines relied on β-oxidation to support OXPHOS (Figure 3c). The etomoxir effect on romidepsin-treated cells was significantly reduced compared with untreated cells for seven out of eight cell lines, indicating that romidepsin inhibits β-oxidation-supported OXPHOS (Figures 3c and Supplementary 3).

**Figure 3:**
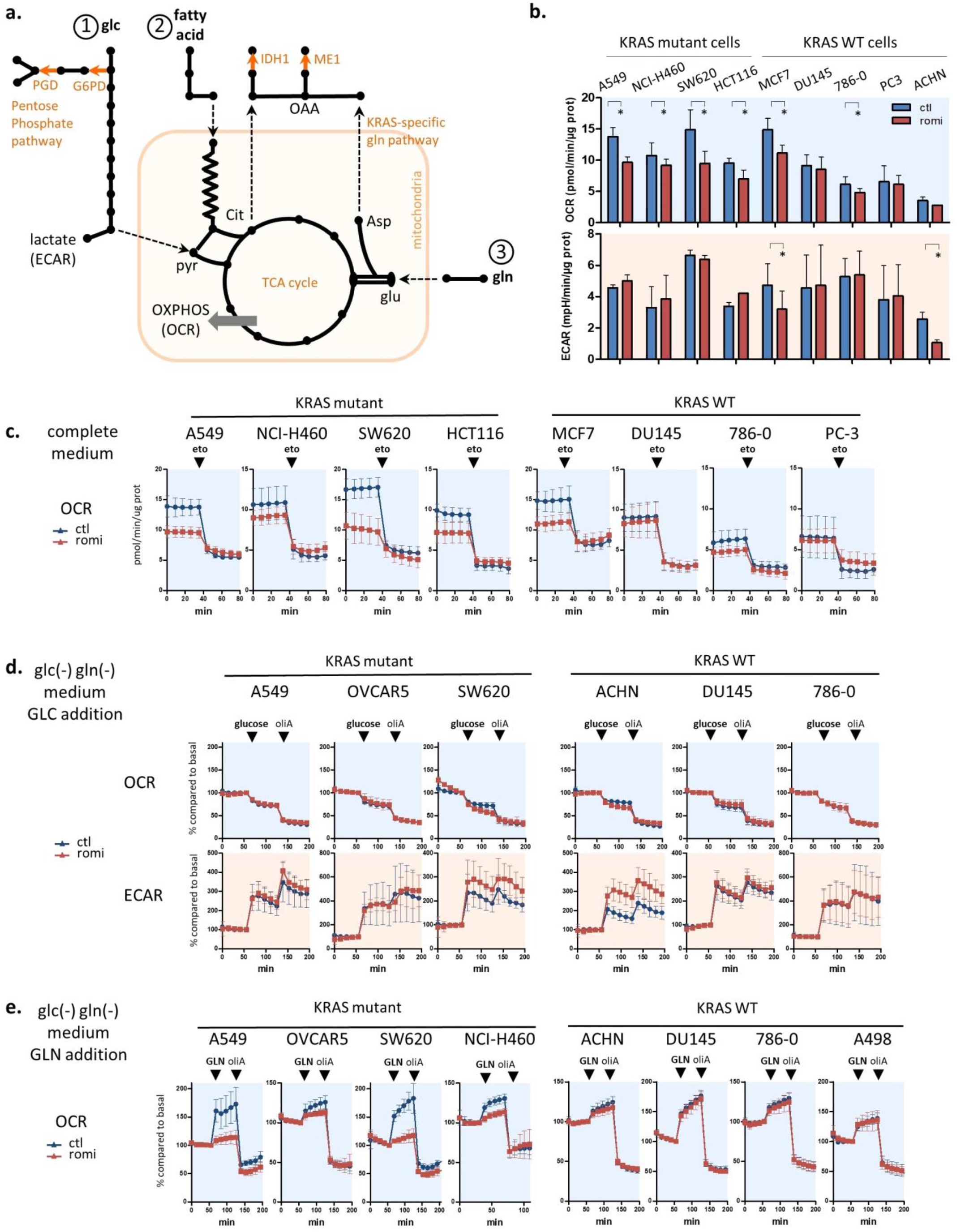
Romidepsin affects two TCA-fueling pathways: fatty acid beta-oxidation and glutaminolysis. **a**. Schematic view of cytoplasmic ROS scavenging through NADPH producing-metabolic enzymes and their link with the three main TCA cycle fueling pathways (1/glycolysis, 2/beta-oxidation, 3/glutaminolysis). Cytoplasmic NADPH producing enzymes are downstream of glutaminolysis in KRAS-mutant cells (ME1), downstream of TCA cycle through citrate production (IDH1), or downstream of the first step of glycolysis (PGD and G6PD in pentose phosphate pathway). **b**. Basal OXPHOS and glycolysis were evaluated by measuring oxygen consumption rate (OCR) and extracellular acidification rate (ECAR) using Seahorse XF96 in four KRAS-mutant and five KRAS-WT cell lines after 18h treatment romidepsin (clinical dosing). **c**. The involvement of fatty acid β-oxidation in TCA cycle fueling was evaluated by adding 40µM etomoxir to cell lines while OCR was measured in four KRAS-mutant and four KRAS-WT cell lines treated for 18h with romidepsin. **d**. The involvement of glycolysis in TCA cycle fueling was evaluated by adding 10mM glucose to glucose/glutamine-free medium while OCR and ECAR were measured in three KRAS-mutant and three KRAS-WT cell lines treated for 18h with romidepsin. **e**. The involvement of glutaminolysis in TCA cycle fueling was evaluated by adding 2mM glutamine to glucose/glutamine-free medium while OCR was measured in four KRAS-mutant and four KRAS-WT cell lines treated for 18h with romidepsin. In **b-e**, data were plotted as mean +/-SD, and statistical significance between groups was assessed using unpaired two-tailed Student’s t-test in **b**.

To assess the role of glycolysis, cells were glucose- and glutamine-deprived for 1h, and then glucose was added to medium while OCR and ECAR were measured. Glucose addition increased glycolysis (ECAR) in all six tested cell lines, pretreated or not with romidepsin (Figure 3d). In contrast, OCR decreased after glucose addition, reflecting a shift from OXPHOS to glycolysis as an immediate source of ATP. The subsequent addition of oligomycin A, which blocks ATP-linked respiration and forces cells to rely on glycolysis to produce ATP, allowed measurement of maximum glycolytic capacity, also unchanged after the HDACi treatment.

To assay whether glutamine was a dominant OXPHOS fuel source rather than glucose, the same experiment was performed in 1h-glucose/glutamine-deprived cells, and glutamine was subsequently added (Figure 3e). In all untreated cells, glutamine addition caused an increase in OCR, indicating that all cells used glutamine for OXPHOS. More importantly, in three out of four KRAS-mutant cell lines, romidepsin inhibited the ability of the cells to shift to OXPHOS after addition of glutamine. This effect was not observed in the four KRAS-WT cell lines.

Together, these results indicated that cancer cell lines used β-oxidation and glutamine rather than glucose as main source of fuel for the TCA cycle, independently of KRAS status. Romidepsin impaired OXPHOS via inhibition of β-oxidation in almost all tested cell lines, but impaired glutamine use for OXPHOS only in cells bearing KRAS mutations.

### Romidepsin affects acetyl-CoA availability

To understand the disturbances in β-oxidation and glutamine metabolism induced by romidepsin, a metabolomic analysis was performed in A549 (KRAS-mutant) and ACHN (WT) cell lines. Cells were treated 12h or 18h with romidepsin using clinical dosing, and 629 metabolites were quantified. Romidepsin-altered pathways were determined by Metabolync pathway enrichment analysis. Fatty acid metabolism and TCA cycle pathways were significantly modified in both cell lines at 12h and/or 18h (Figures 4a and Supplementary 4a), as well as valine/leucine/isoleucine metabolism, another source of the TCA cycle-fueling metabolite acetyl-coA, confirming OCR/ECAR data (Figure 3).

**Figure 4:**
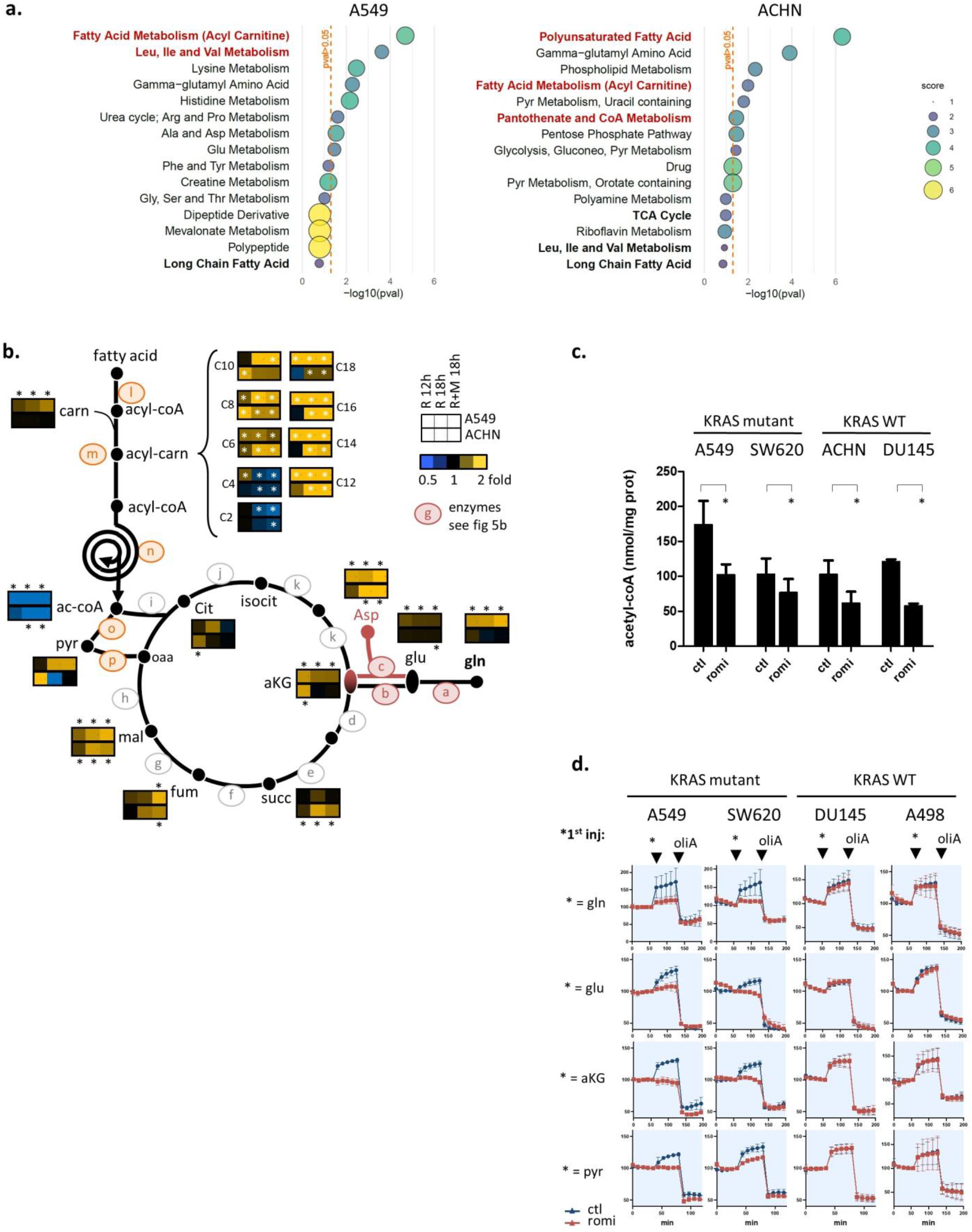
Romidepsin affects acetyl-CoA availability. **a**. 620 metabolites were quantified in A549 and ACHN cell lines after 18h romidepsin treatment (clinical dosing), and the top15 enriched metabolic pathways were obtained after enrichment pathway analysis on significantly modified compounds (Welsh’s t-test). P-values for pathway scores were calculated using hypergeometric test. **b**. 620 metabolites were quantified after treatment with romidepsin (clinical dosing) for 12h, 18h and 18h in combination with MEKi, and those involved in β-oxidation, glutaminolysis and TCA cycle were overlaid on TCA cycle fueling pathway representation (median fold change). Letters refer to enzymes whom expression was quantified by microarray analysis, and for which results are presented in Supplementary Figure 5c. Statistical difference between control and treated groups was evaluated by Welsh’s t-test and asterisk indicated p-values<0.05. **c**. Acetyl-CoA levels were quantified by mass spectrometry in four cells lines following 18h romidepsin treatment. **d**. Cells were treated for 18h with romidepsin, then incubated 1h in medium without glucose and glutamine, and then variation in OXPHOS (OCR) was measured by Seahorse analysis while either glutamine, cell permeant-glutamate, cell permeant-αKG or pyruvate was subsequently added, followed by oligomycin A. In **c-d**, data are plotted as mean +/-SD, and statistical difference was assessed using unpaired two-tailed Student’s t-test.

Metabolites in the TCA cycle and in its two fueling pathways (glutaminolysis and β-oxidation) are reported in Figure 4b, and metabolites from valine/leucine/isoleucine degradation pathway and glycolysis in Supplementary Figure 4b-c. The TCA cycle metabolite malate accumulated in both cell lines after 18h, indicating a block in TCA cycle flux. In the β-oxidation pathway, C6- to C18- acyl-carnitine (upstream) accumulated while acetyl-CoA (downstream) was strongly depleted in both cell lines. These data suggest that romidepsin induced a block in fatty acid flux toward the TCA cycle, similar to what was observed with OCR/ECAR experiments (Figure 3). Glutamine, glutamate and αKG accumulated only in KRAS-mutant cells, confirming as well the decreased ability of these cells to use glutamine for TCA cycle following romidepsin treatment.

Notably, among all the quantified metabolites in the TCA fueling pathway, only acetyl-CoA and its two direct precursors downstream of β-oxidation (C2- and C4-acylcarnitine) were depleted, whereas almost all other TCA-fueling metabolites were accumulated (Figure 4b). To validate the metabolomics analysis, we quantified cellular acetyl-CoA in four cell lines by mass spectrometry. Romidepsin induced a decrease in acetyl-CoA in the two KRAS-mutant cell lines as well as in the two KRAS-WT ones (Figure 4c). Knowing that the rate of flow through TCA is limited by the availability of its substrates oxaloacetate and acetyl-CoA, these data suggest that acetyl-CoA depletion caused a block in TCA cycle metabolite flux in KRAS-mutant cells, consequently causing an accumulation of glutamine downstream metabolites that were feeding into the cycle in the KRAS cells only.

To explore this hypothesis, we returned to the Seahorse experiments. Cells were placed for 1h in a glucose/glutamine-deprived medium, then glutamine, DM-glutamate (cell permeable glutamate), DM-glutarate (cell permeable αKG) or pyruvate were added while OCR was measured (Figure 4d). After 18h romidepsin treatment, KRAS-mutant cells could not use glutamine, glutamate, αKG or pyruvate for OXPHOS, whereas no variation in the use of these metabolites was observed between treated and untreated cells in KRAS-WT cells. The fact that αKG was not used for OXPHOS in romidepsin-treated KRAS-mutant cells indicated that the decrease in glutamine flux induced by romidepsin was not due to a decreased activity in glutaminolysis enzymes (GLS, GOT2 and GLUD1). It indicated an event downstream of the αKG formation step. Romidepsin also inhibited cellular ability to use pyruvate in the KRAS-mutant cells. These findings confirmed that romidepsin induced a block in TCA cycle flux downstream of glycolysis and glutaminolysis, leading to accumulation of unused fueling metabolites (seen in Figure 4b), and a decrease of OXPHOS in KRAS-mutant cells (seen in Figure 3b).

Taken together, we postulated that that romidepsin-induced acetyl-CoA depletion was the cause of TCA cycle flux inhibition.

### Acetyl-CoA precursors replenish acetyl-CoA stock and rescue KRAS-mutant cells from metabolic stress

Acetyl-CoA is a central metabolite that links cell energy status and transcription regulation. It is a TCA cycle limiting factor, as well as the only substrate for histone acetylation, the epigenetic mechanism regulating transcription (22). To gain further insight into romidepsin-modified acetyl-CoA metabolism, microarray analysis was performed on A549, Miapaca2 and ACHN cell lines after 6h or 18h romidepsin treatment (clinical dosing), and pathway enrichment analysis using the KEGG signature database to cover more thoroughly modifications in metabolic pathways. Pathway mining highlighted modifications in fatty acid metabolism, valine/leucine/isoleucine degradation, and steroid biosynthesis in the top10-modified pathways for the three tested cell lines (Supplementary Figure 5a). Nevertheless, even if these pathways are involved in acetyl-CoA metabolism, variation in the expression of acetyl-CoA metabolism genes was not involved in acetyl-CoA depletion, and even trended to counter it with increasing levels of enzymes producing acetyl-CoA and decreasing levels of enzymes using acetyl-CoA (Supplementary Figure 5b-c).

Since modification in metabolic enzyme mRNA level was not the cause of acetyl-CoA depletion, we looked at other possible reasons. Noticeably, acetyl-CoA is the sole acetyl-group donor for protein acetylation. Yet, romidepsin is a histone deacetylase inhibitor, so it causes abnormally high cell acetylation levels, and thus an abnormally high acetyl group consumption that could be the main cause of acetyl-CoA depletion. We therefore assessed whether restoring acetyl-CoA metabolite levels could be sufficient to rescue the cell damages that we observed after romidepsin treatment.

To reintroduce acetyl-CoA in cells, we incubated cells with two cytoplasmic acetyl-CoA precursors (citrate and acetate), because acetyl-CoA itself cannot cross the cytoplasmic membrane. Various precursor concentrations were tested on three KRAS-mutant cell lines and we selected the optimal conditions for cell death rescue (Supplementary Figure 6). Addition of 5mM acetate plus 20mM citrate in romidepsin-treated A549 cell line rescued the KRAS-mutant cells from romidepsin-induced cell death (Figure 5a). We then verified that citrate/acetate did replenish acetyl-CoA stock at this concentration. Mass spectrometry quantification of acetyl-CoA level in A549 and ACHN confirmed that addition of acetate/citrate in romidepsin-treated cells replenished acetyl-CoA stock with levels equivalent to those observed in untreated cells (Figure 5b). These results also indicated that the acetyl-CoA producing enzymes ACLY (catalyzing citrate to acetyl-CoA), ACSS1 (catalyzing acetate to mitochondrial acetyl-CoA) and ACSS2 (catalyzing acetate to cytoplasmic acetyl-CoA) were not themselves affected by romidepsin treatment, neither in terms of transcription (confirming Supplementary Figure 5 results), translation, or post-translation.

**Figure 5:**
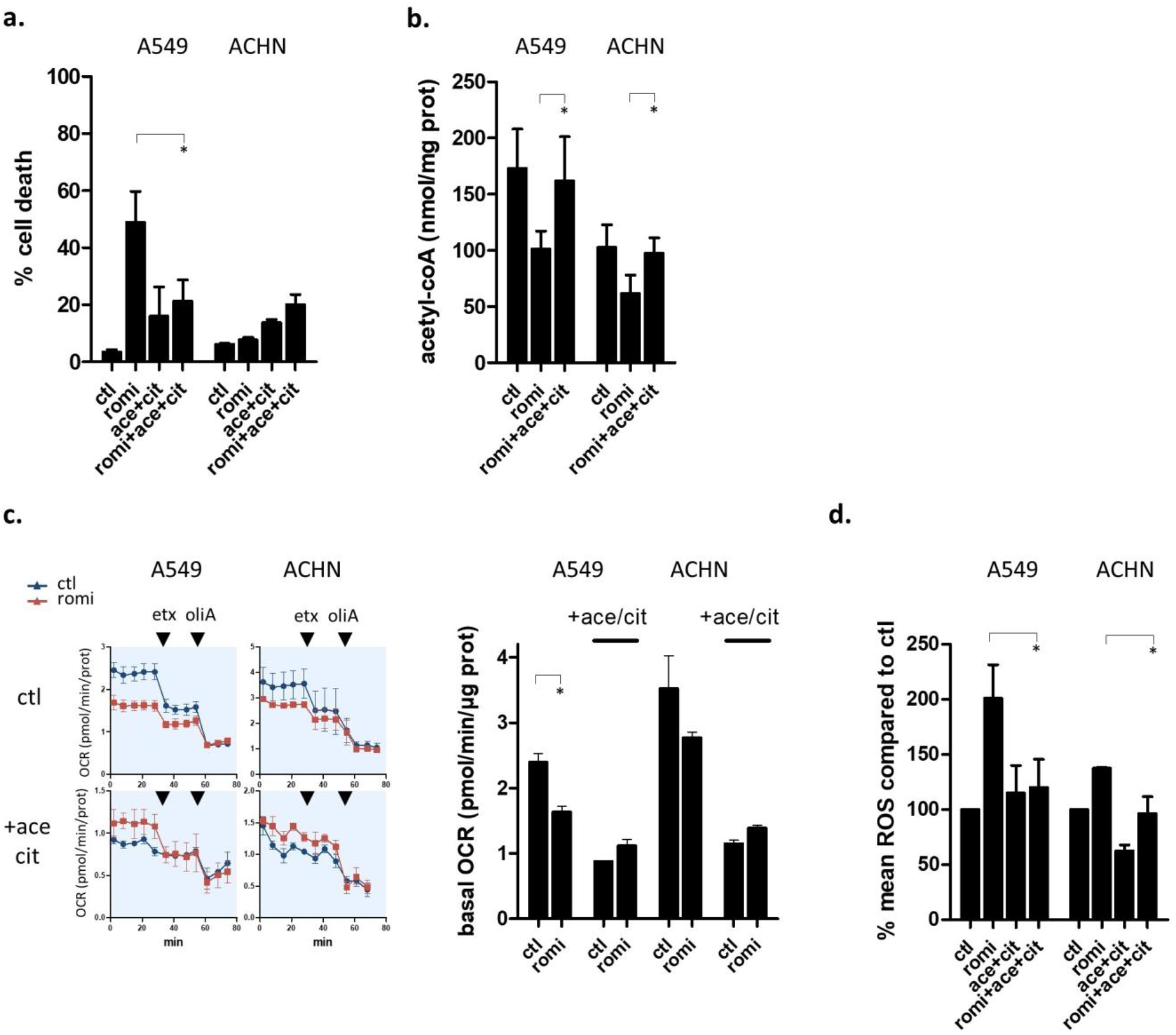
Acetyl-CoA precursors replenish acetyl-CoA stock and rescue KRAS-mutant cells from metabolic stress. **a**. Cell death was measured by flow cytometry (annexin-V positive cells) in cells treated or not for 48h with romidepsin (clinical dosing), in the presence or not of 20mM acetate + 5mM citrate. **b**. Acetyl-CoA was quantified by mass spectrometry in KRAS-mutant (A549) and KRAS-WT (ACHN) cell lines treated or not with romidepsin (18h), in presence or not of 20mM acetate + 5mM citrate. **c**. OXPHOS (OCR) was measured in cells treated or not for 18h with romidepsin, in presence or not of 20mM acetate + 5mM citrate followed by addition of 40µM etomoxir then 5µM oligomycin A (left). Basal OCR level for romidepsin-treated and untreated cells was calculated and plotted according to presence or not of acetyl-CoA precursors in medium (right). **d**. ROS production was measured by flow cytometry (DCF-DA) in cells treated or not for 24h with romidepsin, in presence or not of 20mM acetate + 5mM citrate. For all graphs, data are plotted as mean +/-SD, and statistical difference was assessed using unpaired two-tailed Student’s t-test.

We then investigated the coupling between acetyl-CoA depletion and cell death. In presence of acetyl-coA precursors, romidepsin did not induce a decrease in basal OCR levels in the KRAS-mutant cells, contrary to what was observed in absence of citrate/acetate (Figure 5c). We also measured romidepsin-induced cytoplasmic ROS production in presence or not of acetyl-CoA precursors, and we observed that acetyl-CoA replenishment inhibited romidepsin-induced oxidative stress in KRAS-mutant cells (Figure 5d).

These results showed that the decrease in acetyl-CoA levels was a leading cause of romidepsin-induced cell death in KRAS-mutant cells. Romidepsin-induced acetyl-CoA decrease caused a reduction in TCA cycle flux and an increase in cytoplasmic ROS production that triggered cell death in KRAS-mutant cells only. Notably, romidepsin-induced damage was reversed by just replenishing acetyl-CoA stocks and were not caused by romidepsin-induced inhibition of acetyl-CoA generation pathways, suggesting that it was hyperacetylation-induced acetyl-CoA overconsumption that led to the decrease in acetyl-CoA.

### Two acetyl-CoA metabolism pathways are markers of HDAC inhibitor sensitivity

Because romidepsin-induced acetyl-CoA decrease was observed in all tested cell lines, whereas only KRAS-mutant cell lines showed a decrease in acetyl-CoA-linked cell death after HDAC inhibition, we hypothesized that KRAS-mutant cells were more acetyl-CoA-dependent than KRAS-WT cell lines.

To determine whether HDACi-sensitive cells had an acetyl-CoA altered metabolism at baseline, we decided to derive an HDACi-sensitivity metabolic signature from transcriptomic datasets. We used the Cancer Cell Line Encyclopedia panel (CCLE, 608 cell lines) with known response to eight HDACis to gather enough samples to gain a sufficient statistical power (19). A machine learning approach was taken to identify key metabolic pathways related to HDACi sensitivity; the pipeline and results are illustrated in Supplementary Figure 7. To validate the importance of metabolism for HDACi sensitivity in clinical samples, we selected the significant metabolic pathways obtained from CCLE analysis and analyzed their baseline enrichment in samples obtained from 13 acute myeloid leukemia (AML) patients for whom microarray data prior to treatment and clinical response to belinostat are publically available. Four out of five acetyl-CoA-related pathways were significantly enriched in patients with good clinical response to belinostat (Figure 6a). The leading-edge genes for the five pathways in both CCLE panel and AML patients are overlaid onto pathway representations with gene expression direction in Figure 6b-c. Genes encoding for fatty acid biosynthesis (consuming acetyl-CoA) were overexpressed in HDACi sensitive samples in patients with AML, and in the CCLE panel. Genes encoding for fatty acid β-oxidation (generating acetyl-CoA) were repressed in CCLE HDACi sensitive cells. Genes encoding for valine/leucine/isoleucine degradation and propionyl-CoA catabolism (both consuming and generating acetyl-CoA) were also overexpressed in patients and in CCLE cell lines showing HDACi sensitivity. These results confirmed that samples with sensitivity to HDACi have an altered expression profile for acetyl-CoA metabolism at baseline.

**Figure 6:**
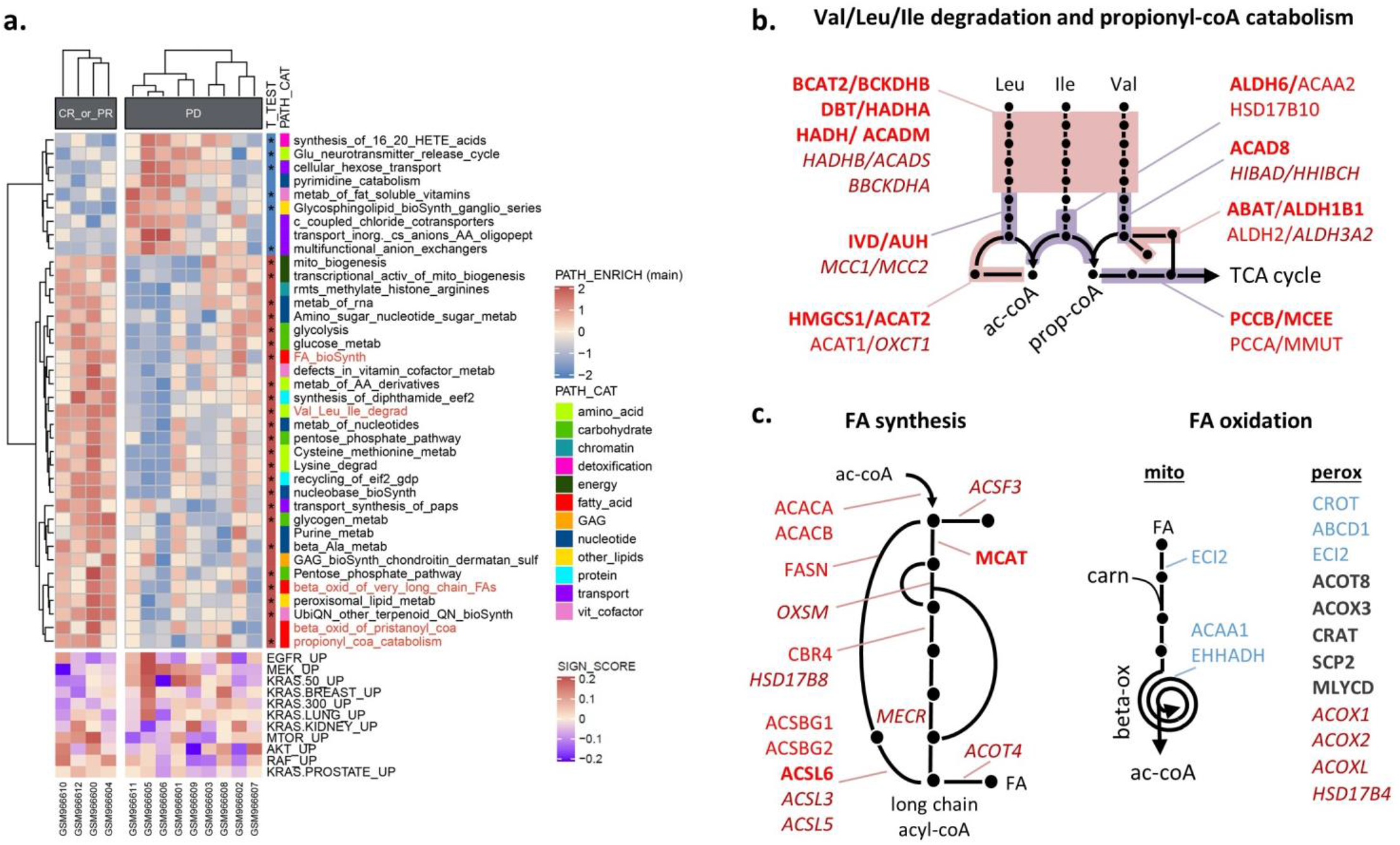
Two acetyl-CoA metabolism pathways are markers of HDAC inhibitor sensitivity. **a**. Key metabolic pathways involved in response to eight HDAC inhibitors were selected by elastic net analysis in CCLE panel (see supplementary Figure 7). The enrichment pathway scores for these pathways were then calculated for 13 AML patients, and plotted according to clinical response to belinostat. RAS pathway activation is indicated below the heatmap while metabolic pathway subcategory and T-test results (* for p-value<0.05) are annotated on the right. The five pathways in red are directly involved in acetyl-CoA generation or consumption. **b. and c**. Leading-edge genes for the five acetyl-CoA-related pathways in AML patients and CCLE panel were overlaid onto acetyl-CoA metabolism pathway representations. Genes in red represent overexpression, in blue: repression and in grey: contradictory results between the two datasets. Bold text indicates that genes were found in both datasets, italic text: in AML patients only, and regular text: in CCLE panel only.

Together, these results demonstrated that the acetyl-CoA metabolism pathway was linked to HDACi sensitivity, and we confirmed that alteration in two acetyl-CoA metabolism pathways (fatty acid metabolism and valine/leucine/isoleucine metabolism) were a hallmark of sensitivity to HDACi in patients.

## DISCUSSION

In this study, we discovered that the class I HDAC inhibitor romidepsin caused a depletion in acetyl-CoA, most probably through an overconsumption of acetyl-CoA itself triggered by induced hyperacetylation of histones. We also uncovered a strategy for treating KRAS pathway modified tumors. Indeed, in RAS-activated cells, the depletion in acetyl-CoA triggered a decrease in TCA cycle flux combined with an increase in cytoplasmic ROS that led to cell death. Computational analysis of transcriptomic profiles in a 608-cell panel highlighted inherent alterations in acetyl-CoA generation/consumption pathways that could cause a higher dependency on acetyl-CoA depleting drugs, and we confirmed our results in patient samples following belinostat. Together, these findings provide a promising strategy for improving treatments for the RAS pathway-modified tumors, where acetyl-CoA generation/consumption signature could be exploited to select patients who are most likely to benefit from HDACi treatment.

The tight link between histone acetylation and acetyl-CoA levels is well known but has been described in only one direction: acetyl-CoA levels determine histone acetylation levels. In fact, the abundance of the nucleo-cytosolic pool has a direct impact on the enzymatic activity of HATs (23). Here, we reported the other direction of this relationship: forcing cells to acetylate histones by HDAC inhibition leads to a depletion in acetyl-CoA. This is consistent with the fact that acetyl-CoA is the unique donor of acetyl groups for acetylation, and that even after 6h treatment followed by 12h washout, the acetylated histone signal did not fade (Supplementary Figure 8), implying that sequestrating the “acetyl group” pool on histones can be a long-lasting event, preventing the quick restoration of acetyl-CoA level. In agreement, Kurdistani (24) hypothesized that acetylated histones might function as a storage site for acetate to be released by HDACs when the need arises. His theory supports our results implying that HDAC inhibition leads to an imbalance in acetyl-CoA pool by inhibiting its release.

In normal physiology, fluctuations in acetyl-CoA concentration reflect the metabolic state of the cell. Due to this key role, its regulation is tightly controlled by three different mechanisms: the expression of its metabolic enzymes or their regulators, its cellular and subcellular compartmentalization, and the post-translational regulation of the enzymes involved in its production (by acetylation) (22,23,25). Several lines of evidence lead us to conclude that it is histone acetylation *per se* rather than any other mechanism that causes depletion of the acetyl pools. First, we evaluated transcription modulation (Figure 5). Second, we indirectly evaluated whether increased protein acetylation could have caused the metabolic defects. Admittedly, ACSS1, ACSS2, ACLY and PDHA1 are involved in acetyl-CoA formation and their activity is inhibited by acetylation (25,26). However, the addition of acetate plus citrate in romidepsin-treated cells restored acetyl-CoA levels to those observed in untreated cells (Figure 6a), indicating that these enzymes were not inhibited. Further, the malate and succinate dehydrogenase (MDH and SDH) enzymes in the TCA cycle are also regulated by acetylation, but OCR/ECAR experiments did not show inhibition of these enzymatic steps (Figure 4c).

Several teams already highlighted the connection between HDACis and energetic metabolism. For example, OCR decrease was induced with pan-HDACis, like butyrate, panobinostat or trichostatin A (27–29). Others have demonstrated that HDACis perturbed fatty acid metabolism, with variation in intermediate metabolites very similar to the ones we observed in our metabolomics analysis (Figure 4b, (30–34)). Although others have shown that HDACis inhibit glycolysis (32,35), there was no significant variation in our hand. These differences could reflect the very important fact that the source of carbon used to fuel metabolism is context dependent. Several teams have indeed demonstrated that glutamine is the predominant carbon source for mitochondrial metabolism *in vitro*, whereas glucose contributes to a greater degree *in vivo* (11,36). Confirming these analyses, in our *in vitro* experiments, cells did not use glucose for lactate production or for OXPHOS, except when they were glutamine-starved, or when beta-oxidation was inhibited. These differences between *in vivo* and *in vitro* experiments could be a barrier to a translation into clinical therapy. Nevertheless, romidepsin-induced tumor growth inhibition in our mouse model, confirming the activity of romidepsin in a KRAS-mutated tumor model in whom glucose is preferentially used over glutamine. Moreover, gene expression analysis from romidepsin-treated cutaneous T-cell lymphoma patients also revealed acetyl-CoA metabolism perturbation (Supplementary Figure 5c). Recently, pancreatic cancer has been shown to undergo reprogramming in lipid-related and acetyl-CoA metabolism pathways (37–39), extending our finding about the importance of acetyl-CoA metabolism in romidepsin sensitivity in KRAS-activated cell lines. Accordingly, a phase II study in patients with KRAS-mutant NSCLC is ongoing with the fatty acid synthase inhibitor TVB-2640 (NCT03808558). Preliminary data indicate pharmacodynamic effects and evidence of clinical activity for TVB-2640 in patients with tumors bearing KRAS mutations (40). These preliminary clinical results are very encouraging, in a field where only a handful of metabolism-targeting drugs have made it into clinical trials, despite the evidence of tumor metabolic high demand.

To conclude, one possible implication of our data is that one of the dominant effect of HDAC inhibition is the depletion of acetyl-CoA. While the canonical mechanism of opening chromatin to facilitate gene transcription is undoubtedly important in some settings, the rapid cell death in T-cell lymphoma cells is more consistent with a metabolic impact. One future direction will be to determine what role acetyl-coA could play in the myriad other effects observed following HDAC inhibition, including R-loop persistence, kinetochore assembly and DNA damage.

## ACKNOWLEDGMENT

This work was supported by the Intramural Research Program (IRP) of the National Cancer Institute (AB/CS/RWR/WML/SEB), the IRP of the National Institute of Diabetes and Digestive and Kidney Diseases (PCV/ML), and the IRP of the Eunice Kennedy Shriver National Institute of Child Health and Human Development (DLS). AB received a fellowship from NIH Intramural Visiting Fellow Program, and funding from the European Union’s Horizon 2020 research and innovation program under the Marie Skłodowska-Curie grant agreement No. 841313. Dr. Bates’ work is supported by a Translational Research grant from the Pancreatic Cancer Action Network and by the Pancreas Center, Columbia University Irving Medical Center. The funders had no role in study design, data collection and analysis, decision to publish, or preparation of the manuscript. The authors thank Dr Tito Fojo for advice over many phases of this project, and the bioinformatics core facility of Nantes (BiRD) for computing resources and support.

## Supplementary Figures

Supplementary Figure 1. Additional information related to Figure 1

Supplementary Figure 2. Additional information related to Figure 2

Supplementary Figure 3. Additional information related to Figure 3

Supplementary Figure 4. Additional information related to Figure 4

Supplementary Figure 5. Variation in acetyl-CoA metabolism gene expression counters romidepsin-induced perturbation

Supplementary Figure 6. Additional information related to Figure 5

Supplementary Figure 7. Analysis of HDACi sensitivity in CCLE panel

Supplementary Figure 8. Histone acetylation over time in cell lines treated with romidepsin (clinical dosing)

