## Supplementary figures for "Histone deacetylase inhibitor induces acetyl-CoA depletion leading to lethal metabolic stress in RAS-pathway activated cells"

**a.**

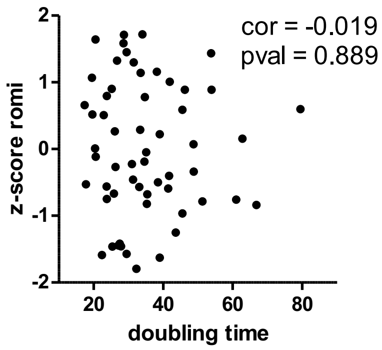


**b.**

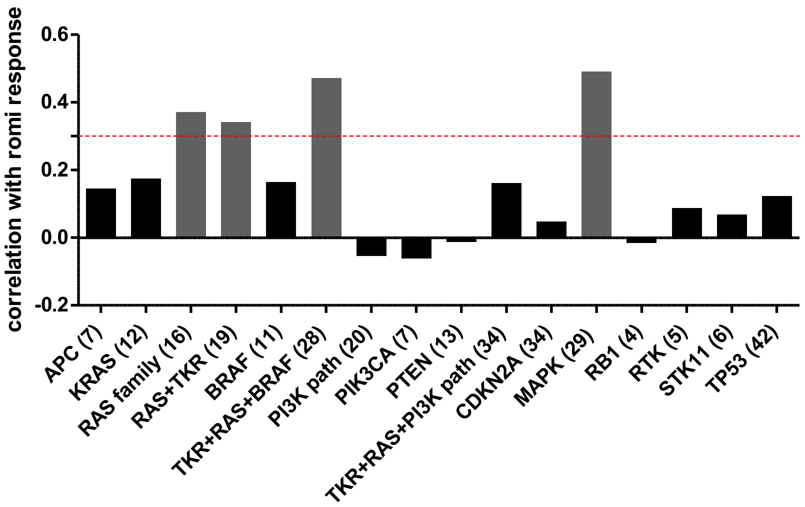


**c.**

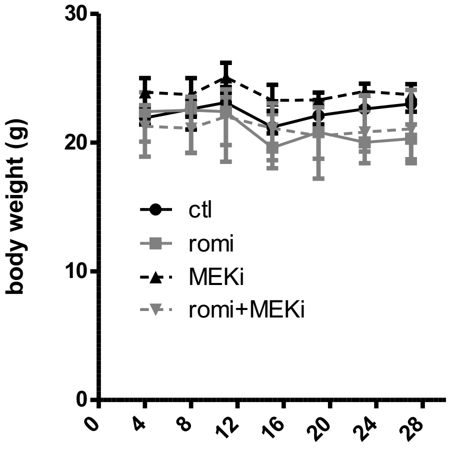


**Supplementary Figure 1 : Additional information related to Figure 1**

**a.** Doubling times for NCI-60 panel (from https://dtp.cancer.gov/discovery_development/nci-60/cell_list.htm) were plotted according to response to romidepsin alone (z-score). **b.** Mutations in key cancer signaling pathways in NCI-60 panel were collected, and their correlation with response to romidepsin was calculated. Correlation analysis was also performed on mutations grouped by pathway, when possible. The number of cells carrying mutation(s) are indicated in parenthesis in x-axis. **c.** Athymic mice were fed with standard mouse chow ad libitum and maintained on a 12h light cycle in pathogen-free conditions. Tumors were generated by injecting subcutaneously 10 million A549 cells resuspended in normal saline into the left flank. Tumor size was measured using a caliper every 4 days, and the average tumor volume ($mm^3$) was calculated as: tumor length × (tumor width)$^2$ x 0.5. Treatment started once tumor size reached an average of 100mm3. In order to achieve a relatively homogeneous distribution between groups, the mice were selected for each treatment group based on approximately equivalent mean tumor size per group on day 24 after cell injection. Treatment started on day 28 after cell injection. Mice received 3mg/kg romidepsin by intraperitoneal injection every 4 days and/or 5mg/kg PD-0325901 by oral gavage every 4 days. PD-0325901 (1mg/ml) was prepared in 0.5% (w/v) hydroxypropyl methylcellulose (HPMC) + 0.2% (w/v) Tween 80. Romidepsin (5 mg/ml in 80% propylene glycol, 20% dehydrated alcohol) was diluted with normal saline to give a concentration of 600µg/ml before injection. Mice were sacrificed after 28 days treatment. Mice weight were measured over time and plotted accordingly (median+/-IQR).

a.

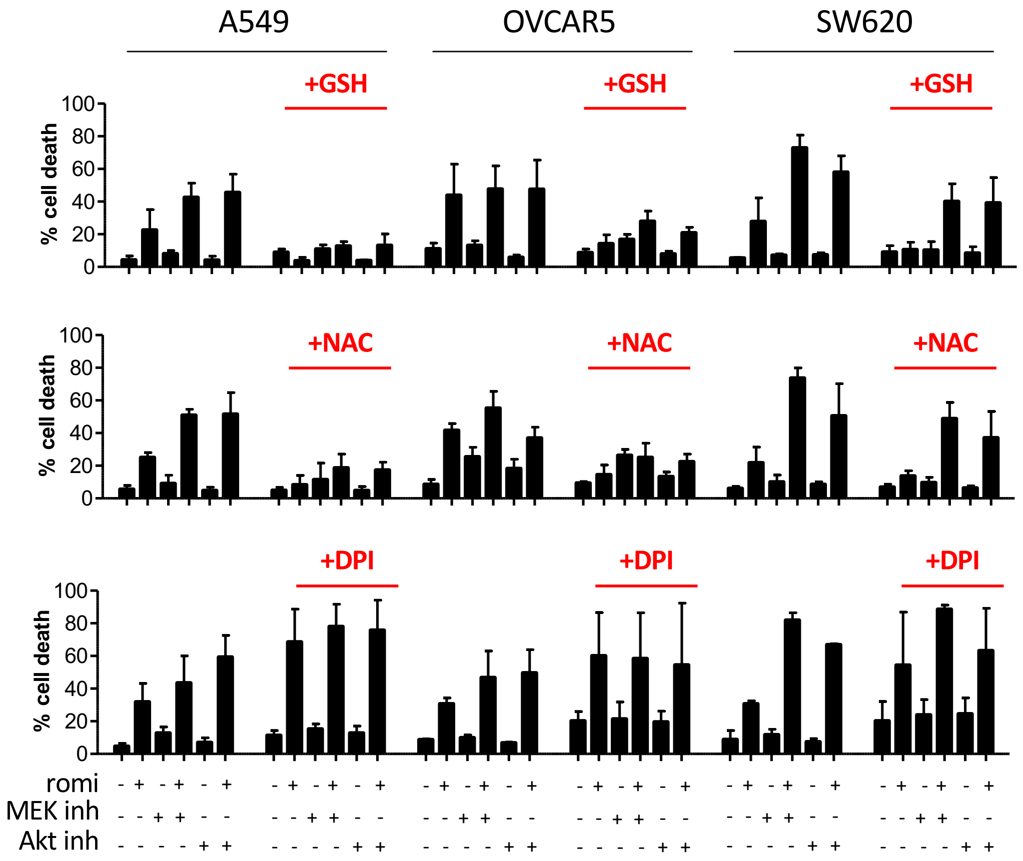


b.

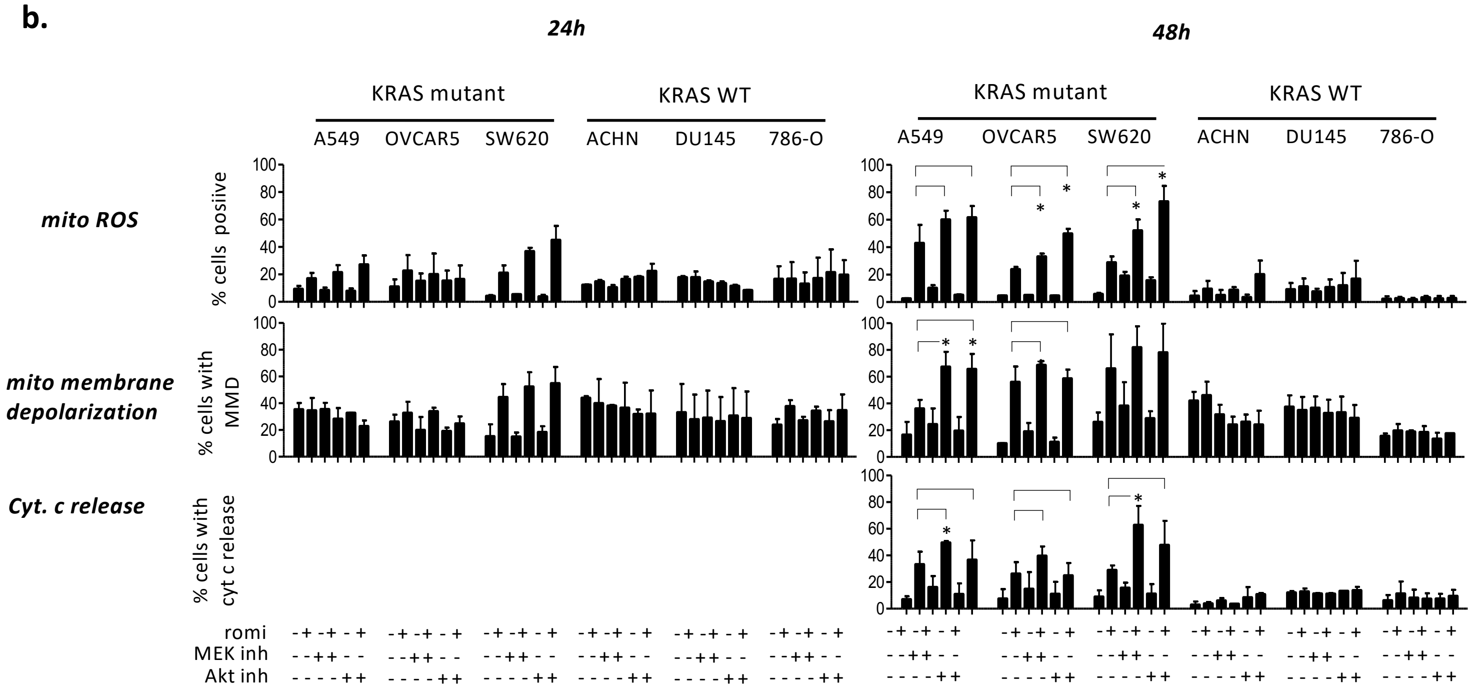


**Supplementary Figure 2 : Additional information related to Figure 2**

**a**. KRAS-mutant cell lines were treated with romidepsin (clinical dosing) +/- MEKi +/- AKTi for 48h, in presence or absence of either ROS scavengers (5mM GSH or NAC) or NOX inhibitors (5µM DPI), then cell death was measured by flow cytometry (annexin-V-FITC). **b.** KRAS-mutant and KRAS-WT cell lines were incubated for 24h and 48h with romidepsin (clinical dosing) +/- MEKi +/- AKTi , then mitochondrial ROS, mitochondrial membrane depolarization and cytochrome c release were measured by flow cytometry (using JC1, MitosoxTM Red and antibody-based cytochrome c release kit respectively). For all graphs, data are plotted as mean +/-SD.

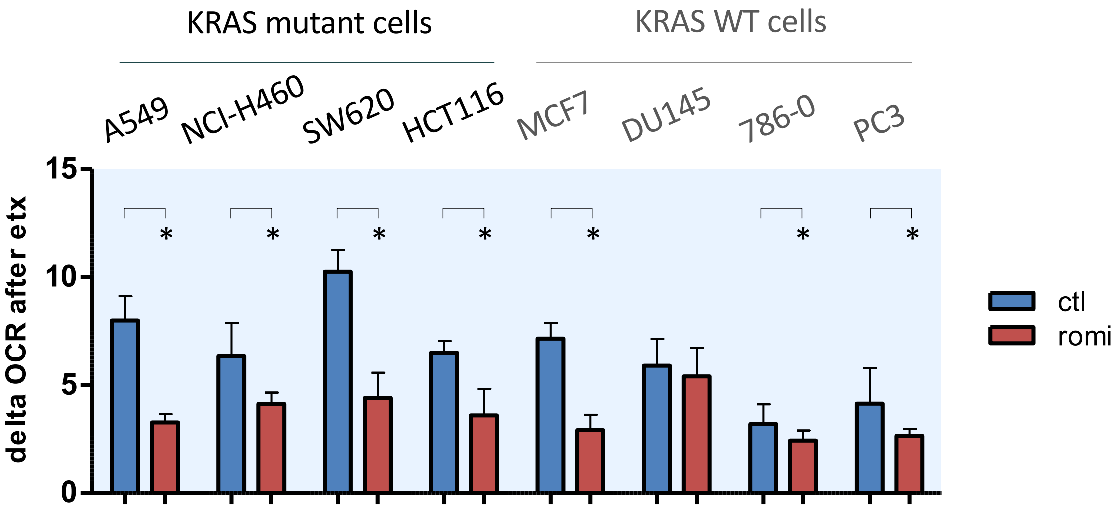


**Supplementary figure 3: Additional information related to Figure 3**

The involvement of fatty acid β-oxidation in TCA cycle fueling was evaluated by adding 40µM etomoxir to cell lines while OCR was measured using Seahorse XF96 in four KRAS-mutant and four KRAS-WT cell lines treated for 18h with romidepsin (clinical dosing). Difference in OCR before and after beta-oxidation inhibition was calculated and plotted accordingly (mean +SD). Statistical significance was assessed using unpaired two-tailed Student's t-test.

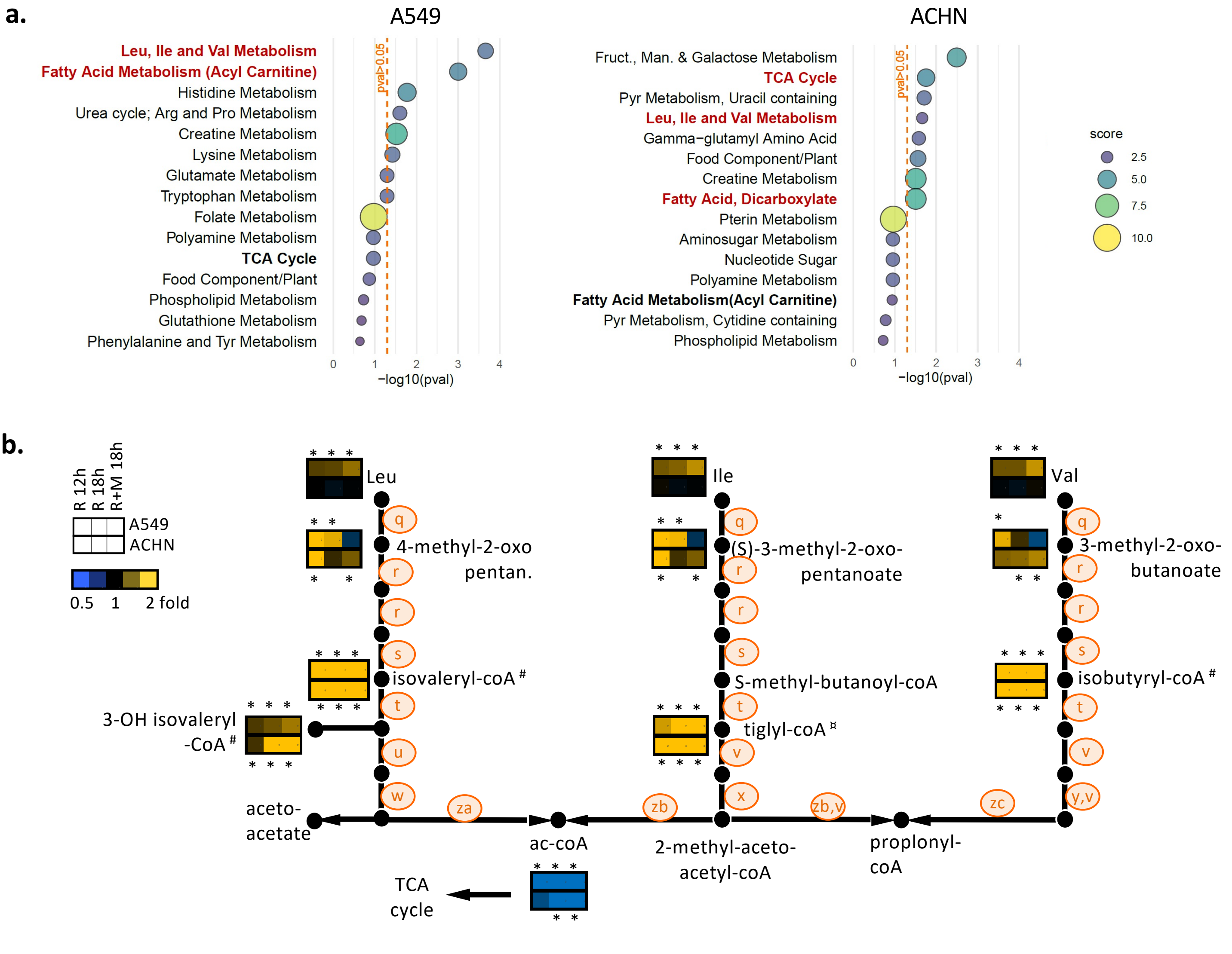


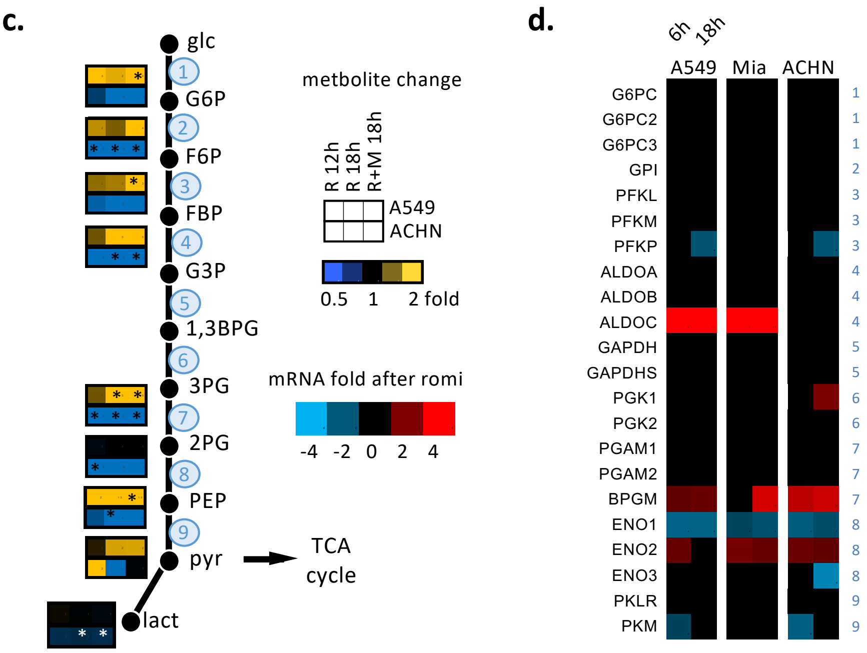


**Supplementary figure 4 : Additional information related to Figure 4**

**a.** 620 metabolites were quantified in A549 and ACHN cell lines after 12h romidepsin treatment (clinical dosing), and the top15 enriched metabolic pathways were obtained after enrichment pathway analysis on significantly modified compounds (Welsh's t-test p-value <0.01). P-values for pathway scores were calculated using hypergeometric test.

**b and c.** 620 metabolites were quantified after treatment with romidepsin (clinical dosing) for 12h, 18h and 18h in combination with MEKi, and those involved in glycolysis (**b**) and valine/leucine/isoleucine degradation (**c**) were overlaid on TCA cycle fueling pathway representation (median fold change). Numbers or letters on schematic view of pathways refer to enzymes whom expression was quantified by microarray analysis, and for which results are presented in panel d, or in Supplementary Figure 5c. Statistical difference between control and treated groups was evaluated by Welsh's t-test and asterisks indicate p-values inferior to 0.05. Hash marks (#) indiquate the quantification of the carnitine form (mitochondrial carnitine shuttle) **d.** All the genes involved in glycolysis were listed and gene expression variations were overlaid onto glycolysis pathway representation. Numbers on the right side of the heatmap refer to enzymatic steps in panel c.

**a.**

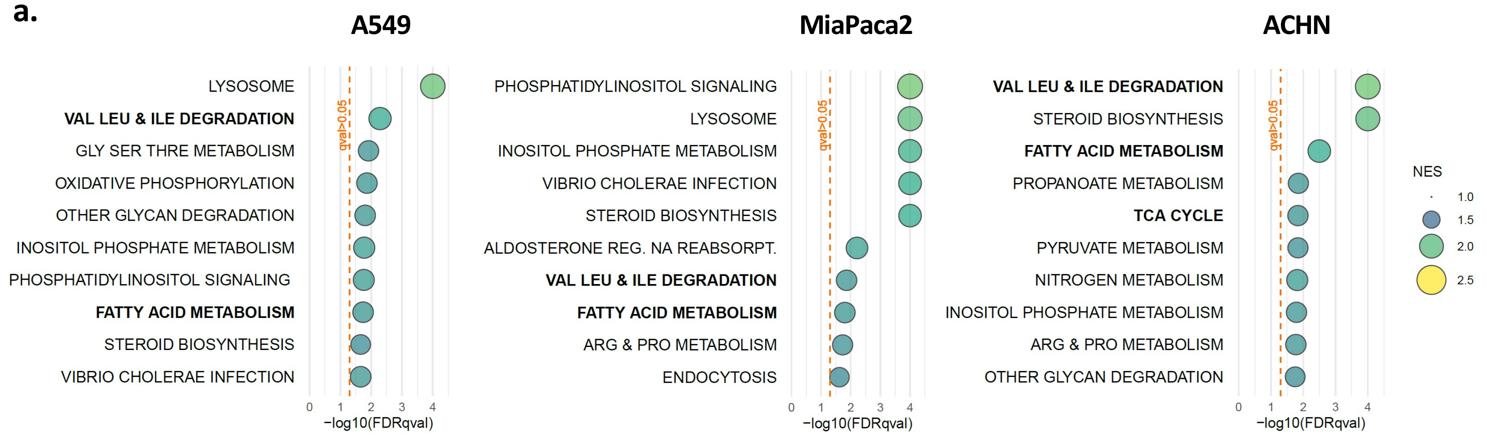


**b.**

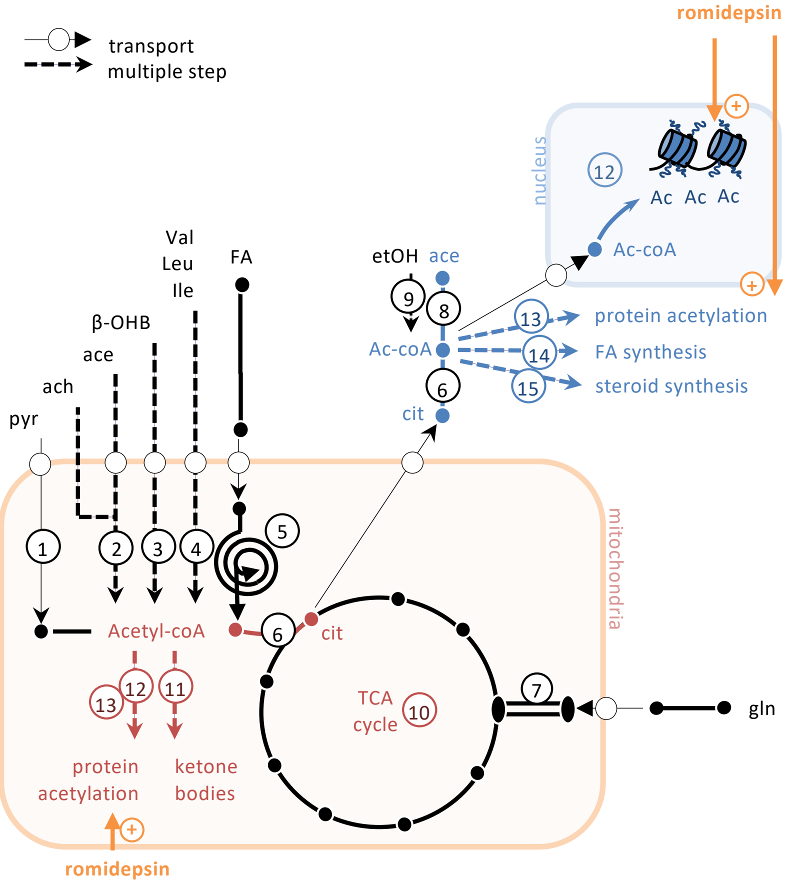


**Supplementary figure 5: Variation in acetyl-CoA metabolism gene expression counters romidepsin-induced perturbation a.** About 17,000 gene transcripts were quantified in A549, MiaPaca2 and ACHN cell lines after 6h and 18h romidepsin treatment (clinical dosing), and the top10 enriched pathways at 18h were obtained after analysis by GSEA using KEGG signature database. **b.** All the genes involved in acetyl-CoA metabolism were listed and overlaid onto acetyl-CoA metabolism pathway representation. The main sources of mitochondrial acetyl-CoA are pyruvate and β-oxidation. The main source of cytoplasmic acetyl-CoA is mitochondrial acetyl-CoA which is translocated into the cytoplasm after forming citrate (which, contrary to acetyl-CoA, is able to cross the mitochondrial border), and is catalyzed back to acetyl-CoA form by cytoplasmic ACLY. Mitochondrial acetyl-CoA is used as substrate for protein acetylation, ketone body formation and is a limiting factor for the TCA cycle. Cytoplasmic acetyl-CoA is a substrate for protein and histone acetylation, fatty acid synthesis and steroid synthesis. **c (next page).** Gene expression variation for the three cell lines and six patients treated with romidepsin are were plotted on heatmaps. Letters on the right side of heatmaps reference to steps in Figure 4b and Supplementary Figure 4b. Gene expression for the patients treated romidepsin were downloaded from GEO (dataset GSE45405). More precisely, peripheral blood mononuclear cells from cutaneous T-cell lymphoma (CTCL) patients were obtained before infusion, at 4h and at 24h after the start of infusion of the first cycle of treatment, then samples were hybridized on Illumina WG-8v2 human whole-genome bead arrays and analyzed for cDNA quantification. First, principal component analysis was performed on the downloaded expression matrix to check for the presence of outliers, and patient 7 was excluded from analysis accordingly. Fold changes in expression after 4h and 24h romidepsin treatment were calculated for patients 1 to 6.

c.

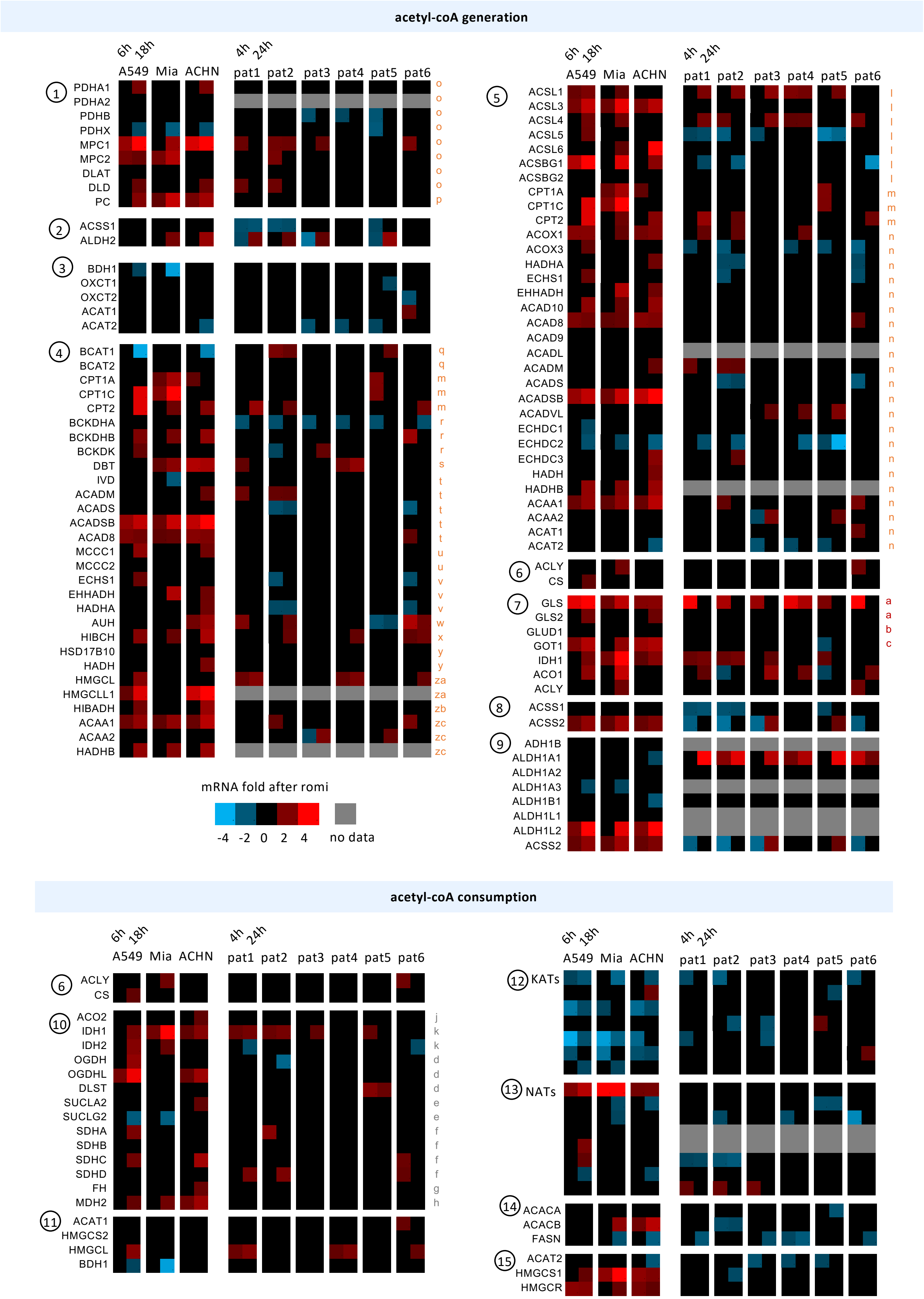

acetyl-coA generation
6h 18h
A549 Mia ACHN
4h 24h
pat1 pat2 pat3 pat4 pat5 pat6
1 PDHA1 PDHA2 PDHB PDHX MPC1 MPC2 DLAT DLD PC
2 ACSS1 ALDH2
3 BDH1 OXCT1 OXCT2 ACAT1 ACAT2
4 BCAT1 BCAT2 CPT1A CPT1C CPT2 BCKDHA BCKDHB BCKDK DBT IVD ACADM ACADS ACADSB ACAD8 MCCC1 MCCC2 ECHS1 EHHADH HADHA AUH HIBCH HSD17B10 HADH HMGCL HMGCLL1 HIBADH ACAA1 ACAA2 HADHB
5 ACSL1 ACSL3 ACSL4 ACSL5 ACSL6 ACSBG1 ACSBG2 CPT1A CPT1C CPT2 ACOX1 ACOX3 HADHA ECHS1 EHHADH ACAD10 ACAD8 ACAD9 ACADL ACADM ACADS ACADSB ACADVL ECHDC1 ECHDC2 ECHDC3 HADH HADHB ACAA1 ACAA2 ACAT1 ACAT2
6 ACLY CS
7 GLS GLS2 GLUD1 GOT1 IDH1 ACO1 ACLY
8 ACSS1 ACSS2
9 ADH1B ALDH1A1 ALDH1A2 ALDH1A3 ALDH1B1 ALDH1L1 ALDH1L2 ACSS2
mRNA fold after romi
-4 -2 0 2 4 no data
acetyl-coA consumption
6 ACLY CS
10 ACO2 IDH1 IDH2 OGDH OGDHL DLST SUCLA2 SUCLG2 SDHA SDHB SDHC SDHD FH MDH2
11 ACAT1 HMGCS2 HMGCL BDH1
12 KATs
13 NATs
14 ACACA ACACB FASN
15 ACAT2 HMGCS1 HMGCR

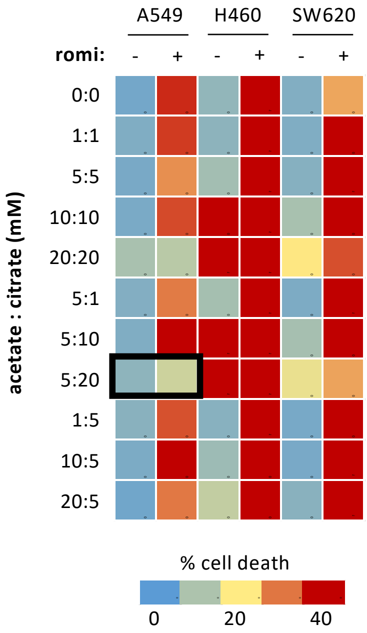


**Supplementary figure 6 : Additional information related to Figure 5**

Optimal acetate/citrate concentration were selected for cell death rescue experiment. Three KRAS-mutant cell lines were treated or not with romidepsin (clinical dosing), in presence or not of various concentrations of citrate plus acetate. Cell death was assessed at 48h by annexin-V-PI assay. Percentage cell death was calculated for each condition and a heatmap was drawn accordingly. The black rectangle indicates the optimal condition.

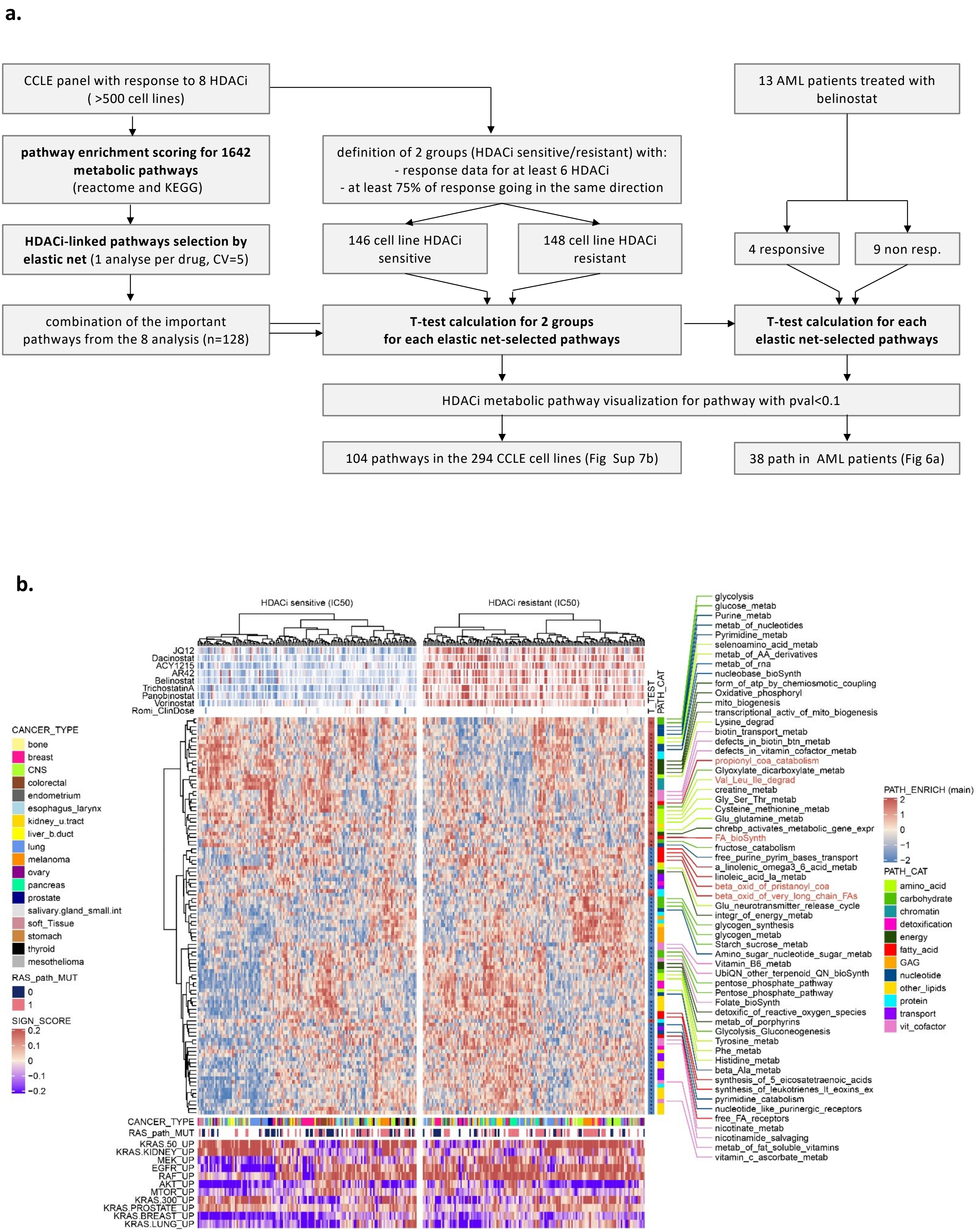


**Supplementary figure 7 : Analysis of HDACi sensitivity in CCLE panel**

See explanation on the next page

**Supplementary figure 7 (previous page): Analysis of HDACi sensitivity in CCLE panel**

**a.** In order to determine metabolic pathways associated with HDACi sensitivity, we performed a machine learning analysis in CCLE panel. First, to reduce number of variables, we calculated enrichment score for 1642 metabolic pathways in the 608 cell lines for which sensibility to at least one HDAC inhibitor was known, using GSVA package. The used gene sets for enrichment analysis were derived from KEGG and Reactome databases (downloaded from Molecular Signatures Database) and have been filtered to remove non-metabolic pathways. Then, for each of the eight HDACi, resistant versus sensible cell lines were split according to median IC50 and an elastic net variable selection was performed on pathway enrichment score according to drug sensitivity class (glmnet package, a = 0.1, lambda= min). A five-time cross-validation (4:1 ratio) was performed by splitting train and test population with respect to class balance (caret package). The 100 first importance-ranked pathways present in at least four on five cross-validation were collected for each of the eight HDACi, and merged. We then selected the cell lines in CCLE panel for which 1/ response to at least 6 on 8 HDACi were known, 2/ at least 75% of response were going in the same direction. In that way, 146 cell lines were selected as HDACi sensitive, and 148 cell lines as HDACi resistant. For each elastic-net selected pathways, we compared enrichment score between the HDACi sensible versus resistant group by T-test, and pathways that had at least a p-value <0.1 were plotted in a heatmap (**b.**) using ComplexHeatmap package. In addition, gene signature scores for 11 RAS signaling pathways were calculated by weighted averaged expression using both up- and down-regulated genes described in C6 collection in Molecular Signatures Database.

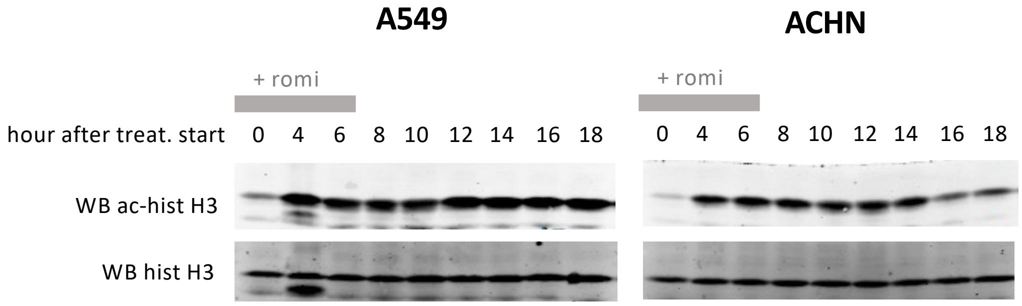


**Supplementary figure 8 : Histone acetylation over time in cell lines treated with romidepsin (clinical dosing)**

KRAS-mutant A549 cells and KRAS-WT ACHN cells were treated for various time point with romidepsin (clinical dosing), then histone acetylation and H2AX phosphorylation were measured by western blot
